# Microstate EEGlab toolbox: An introductory guide

**DOI:** 10.1101/289850

**Authors:** Andreas Trier Poulsen, Andreas Pedroni, Nicolas Langer, Lars Kai Hansen

## Abstract

EEG microstate analysis offers a sparse characterisation of the spatio-temporal features of large-scale brain network activity. However, despite the concept of microstates is straight-forward and offers various quantifications of the EEG signal with a relatively clear neurophysiological interpretation, a few important aspects about the currently applied methods are not readily comprehensible. Here we aim to increase the transparency about the methods to facilitate widespread application and reproducibility of EEG microstate analysis by introducing a new EEGlab toolbox for Matlab. EEGlab and the Microstate toolbox are open source, allowing the user to keep track of all details in every analysis step. The toolbox is specifically designed to facilitate the development of new methods. While the toolbox can be controlled with a graphical user interface (GUI), making it easier for newcomers to take their first steps in exploring the possibilities of microstate analysis, the Matlab framework allows advanced users to create scripts to automatise analysis for multiple subjects to avoid tediously repeating steps for every subject. This manuscript provides an overview of the most commonly applied microstate methods as well as a tutorial consisting of a comprehensive walk-through of the analysis of a small, publicly available dataset.

## 1 Introduction

Multichannel electroencephalography (EEG) is used to assess the spatio-temporal dynamics of the electrophysiological activity of the brain. Traditionally, researchers analyse EEG by characterising the temporal waveform morphology and/or frequency distribution of recordings at certain preselected electrodes. Even though this method provides a wealth of insights into the electrophysiology of the human brain at work and at rest, this approach still misses out on large parts of the information in the EEG signal and neglects the multivariate characteristics of the measurements. A different approach, that takes into account the information of all electrodes, is to characterise the EEG signal by the spatial configuration of the electric fields at the scalp that can be conceptualised as a topographical map of electrical potentials. The widely used EEGLAB toolbox (Delorme and Makeig, 2004) has promoted the use of independent component analysis (ICA) for such multivariate modelling.

*Microstate analysis* is an alternative, and increasingly applied, EEG-representation based on topographic analysis. Microstate analysis has its foundation in the work of Dietrich Lehmann and colleagues (e.g. Lehmann, 1971; Lehmann et al., 1987), who observed that the time series of topographies in ongoing EEG are comprised of a discrete set of a few prototypical topographies that remain stable for around 80 - 120 ms before rapidly transitioning to a different topography. These periods of quasi-stable EEG topography have been called *functional microstates* and the discrete spatial configurations *microstate classes*. Following the argument that different EEG topographies are generated by different configurations of neuronal generators, microstates have been suggested to reflect global functional states of the brain (Khanna et al., 2015; Michel et al., 2009; Lehmann et al., 1998) and have been found to be tightly coupled with functional Magnetic Resonance Imaging (fMRI) resting state networks (Britz et al., 2010; Yuan et al., 2012; Musso et al., 2010; Van de Ville et al., 2010).

Microstate analysis is increasingly recognised as an innovative method offering straight-forward characterisations of brain-states. Its usefulness in gaining insights into brain functions has been demonstrated in healthy (e.g. Koenig et al., 2002) and clinical populations (e.g. Lehmann et al., 2005). Microstates have been examined when subjects are at rest (e.g. Khanna et al., 2015), as well as both during and in between active tasks (e.g. Pedroni et al., 2017). More recently, microstate analysis has been devised in event-related study designs, examining characteristics of pre-and post-stimulus time-locked microstates (e.g. Schiller et al., 2016; Ott et al., 2011).

The output parameters of microstate analysis (described in sections 2.6 and 3.9) quantify the statistical properties of the microstates. They include parameters summarising the variability in the spatial configuration and strength of the topography of microstate classes, parameters pertinent to the temporal dynamics of microstates such as the average duration of a microstate class or the frequency of occurrence, as well as statistics on the transition probabilities between microstate classes. In addition, parsing the EEG into microstates can be used to select epochs of interest (that correspond to a certain microstate class), which can be further examined using other analysis methods such as time-frequency analysis. Also from a practical point of view, compared to functional magnetic resonance imaging (fMRI), microstates analysis of EEG offers a readily applicable and cost-worthy method to investigate temporally coherent network activity.

The core of microstate analysis is to segment the EEG recordings into microstates using a clustering method. There are currently several cluster methods being used for microstate analysis, such as (modified) K-means clustering (Lloyd, 1982; Pascual-Marqui et al., 1995), Topographic Agglomerative Hierarchical clustering (Murray et al., 2008; Khanna et al., 2014), principal component analysis (Skrandies, 1989), or mixture of Gaussian algorithms, with each method reflecting rather a class of clustering algorithms than a completely defined algorithm.

Microstate analysis could arguably have been even more widely adopted had it not been for the relative lack of transparency regarding the methods applied in the field. So far, established toolboxes for microstate analyses have been implemented in compiled software^1^, which therefore not revealing all information of the methods. However, transparency about the exact method is in our view important to grant replicability because, as our experience has shown, even small changes in the settings of clustering algorithms sometimes lead to substantial differences in the outcome of the analysis.

The present work aims to facilitate microstate analysis in Matlab by providing a toolbox that is fully transparent with respect to all the steps of analysis and which allows the integration of any clustering algorithm. The toolbox is made of a set of functions that can be used and modified independently or as an interactive plug-in for the widely used, open-source EEG analysis software EEGLAB (Delorme and Makeig, 2004).

We encourage the reader to give suggestions for improvement or self-developed extensions that may be of interest for a broader community. As the development of clustering algorithms is a rapidly advancing field of research, an open and easily modifiable platform to implement new clustering methods for microstate analysis may also foster new approaches to the idea of EEG microstates. For instance, we see potential in advancing methods to select an adequate number of clusters.

The remainder of this article is structured as follows: Section 2, starts by outlining the general methods to conduct a microstate analysis and then provide detailed explanations about three clustering algorithms (i.e. *K-means, modified K-means* and *(Topographic) Atomize and Agglomerate Hierarchical Clustering*) that have been commonly used in microstate analyses. We then outline some of measures of fit that can be used selecting the number of microstates, which is followed by a discussion on the settings of microstate clustering. We conclude the methodological part of this guide with by explaining the back-fitting procedure and with an overview of the output of a microstate analysis; the microstate statistics. Section 3 is dedicated to provide an easy entry point into microstates analysis in the form of a tutorial that uses the GUI of the toolbox along with a Matlab script, allowing to automatise and adapt the analysis to new datasets.

## 2 Microstate methods

This section contains the theoretical part of the guide, where we will walk through the concepts and differences between the methods employed in microstate analysis. We will cover some of the most common clustering algorithms and their settings, as well as approaches on how to select the best solution and how to evaluate it.

In microstate analysis it is the goal to segment the recorded EEG time samples into microstate classes, so EEG samples that belong to the same class have as similar topographies as possible. The segmentation is commonly done using topographical clustering methods, where each cluster of EEG samples denotes a microstate class. For each cluster, the methods calculate a prototypical topographical map based on all the EEG samples assigned to it. The assumption of the clustering is that EEG samples assigned to the same cluster, all originate from the neural processes underlying the prototype topography of that cluster.

In general there are different mathematical and statistical approaches to cluster data, resulting in different clustering methods. How large a differences there is in the resulting clusters, between the clustering methods, depends on the data of interest, and the underlying assumptions of the methods. For example, where some methods ignore the polarity^2^ of the topographies, other methods take it into account and assign EEG samples with similar but opposite polarity to different clusters. In section 2.1 we go through the most common clustering methods used in microstate analysis.

After deciding on a clustering method there is, however, still factors that affect the clustering, such as; *How to measure the similarity between two EEG samples, should the cluster assignment of a sample be affected by its temporally neighbouring samples*, and *what is the right amount of clusters (microstate classes)*? And after clustering the data it may be relevant to reduce noise by temporal smoothing of the obtained microstate sequence, before calculating statistics from them.

Another thing to note about clustering is that often there is not one true solution that segments the EEG perfectly. In the same way that it is necessary to decide which amount of microstate clusters gives the best result (see section 2.2), some clustering algorithms can converge on different solutions even when using the same number of clusters. This means that using the same method with the same number of clusters might result in different segmentations, when it is run multiple times. See section 2.3.1 for more information on initialisation of algorithms and using ‘multiple restarts’ to reduce variability.

In the following sections we will describe the different methodological choices that can be made in microstate analysis, with the intent of increasing the understanding of the effect of these choices and how they work. For a walk-through of an example microstate analysis we refer to the tutorial in section 3.

During our review of the methodological side of microstate literature, we encountered several examples of citation of articles that were irrelevant for the cited method, or even citation of articles featuring a completely different method. We have therefore made an emphasis on referencing the relevant articles during the review of microstate methods, with the intention that this section can be used to look up the proper references for specific methods.

### 2.1 Clustering algorithms

This section contains a short review of the methods implemented in the Microstate toolbox. The toolbox contains the three most commonly used clustering methods for microstate analysis; *K-means, modified K-means* and *Topographic Atomize and Agglomerate Hierarchical Clustering*. The toolbox also contains a subgroup of new experimental methods under *Experimental algorithms*.

To help explain the methods we introduce the notation in table 1. The same notation is used in the Matlab code of the toolbox. Note that in microstate literature it is in general assumed that the EEG has been referenced to the average reference.

**Table 1:**
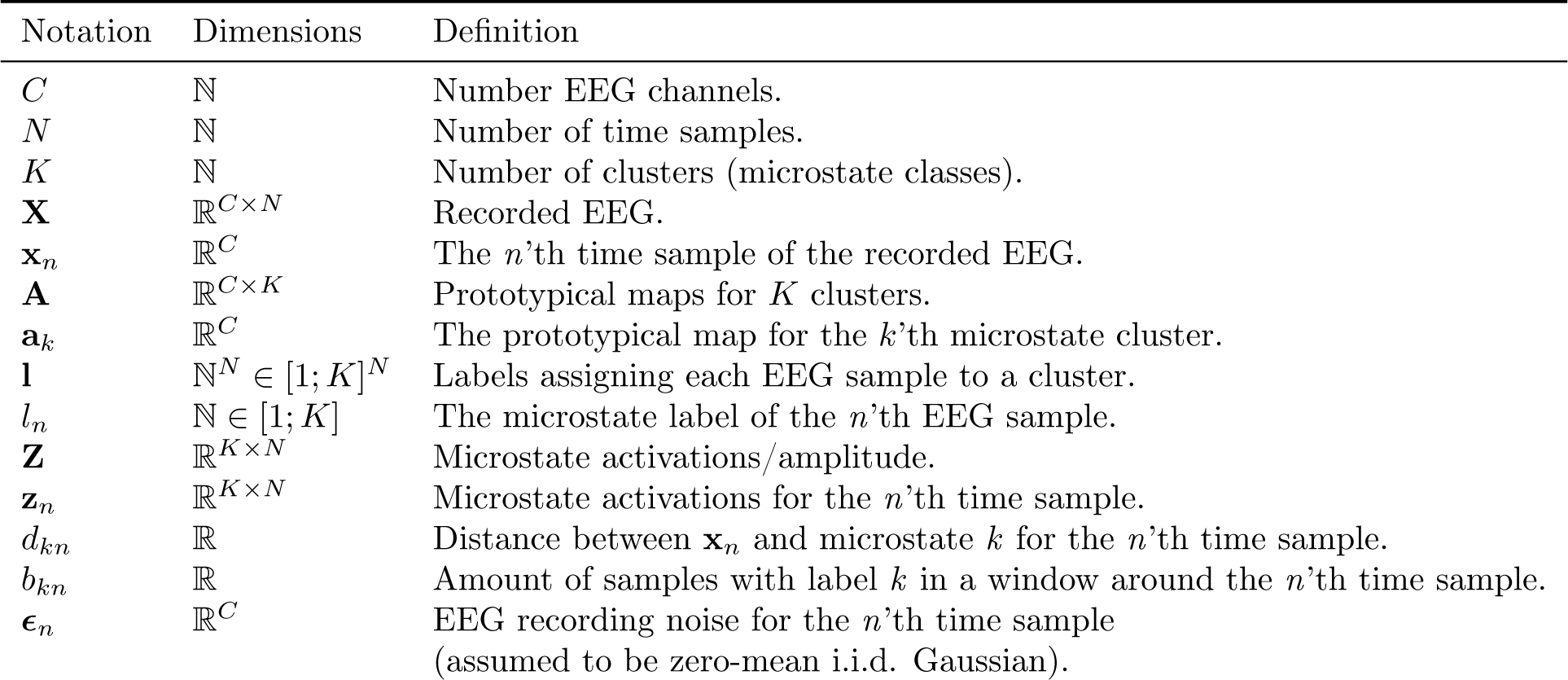
Definitions of the notation employed in this article and in the toolbox.

#### 2.1.1 K-means

K-means clustering (Lloyd, 1982) is a well-established clustering method and also the most basic method employed in the toolbox. It belongs to one of two main groups of clustering methods named the *partitioning methods*, which generally require that the number of clusters (or microstate classes) are pre-set by the user. These methods start with partitioning the EEG samples into the fixed number of clusters, which they relocate the EEG samples to in iterations, until an optimal cluster assignment have been achieved (Rokach and Maimon, 2005).

K-means clustering starts by defining *K* cluster centres (also known as prototypes), e.g. by choosing *K* EEG samples at random. It iterates through two steps: First it assigns each EEG sample to the cluster whose prototype it is most similar with. It then re-calculates cluster prototypes, which is often done by averaging over the newly assigned samples. The algorithm continues iterating over these two steps until a convergence criterion has been reached. Two examples of criteria of convergence could be: Stopping when the change in assignments of EEG samples between iterations are low enough to reach a pre-set threshold or when a fixed number of iterations are reached.

From a probabilistic point of view K-means is consistent with the generative model

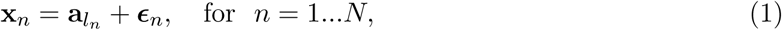

where 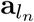 signifies the topographical map assigned to *n* th EEG sample.

K-means clustering can be customised e.g. by changing its way of initialising prototypes or how similarity is measured. The K-means clustering implemented in this toolbox employs Matlab’s default built-in function **kmeans.m** (Statistics and Machine Learning toolbox required), that uses the **k-means**++ algorithm (Arthur and Vassilvitskii, 2007) for cluster prototype initialisation and the squared Euclidean metric to determine similarity.

See e.g. Bishop (2006) or Rokach and Maimon (2005) for a more detailed introduction to K-means clustering.

#### 2.1.2 Modified K-means

The modified K-means introduced in Pascual-Marqui et al. (1995) adds a number of features to clustering.

On a practical level there are mainly two differences compared to conventional K-means. The first is that the topographical maps of the prototypical microstates are polarity invariant. This means that samples with proportional, but opposite, topographical maps (e.g. **a**_*k*_ and −**a**_*k*_) are assigned to the same cluster. The second difference is that modified K-means models the activations of the microstates, i.e. models the strength of the microstates for each time point.

At the conceptual level, microstate classes (or clusters) are now being viewed as ‘directions’ in a multi-dimensional topographical space. The activations would then quantify how far along a microstate-orientation the EEG signal is at a given time point. It makes it easier to imagine this using only two EEG channels, which would draw an asterisk in 2D-space. Here each line is a microstate, and time points with a strong EEG signal would be placed further along its microstate line.

Again, taking a probabilistic view point modified K-means is consistent with the generative model

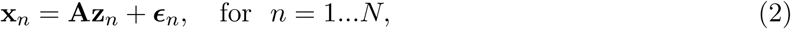

with the important constraint that only one microstate can be active for each time point, i.e. all *K* elements of **z**_*n*_ are zero except for one, i.e. the model can be written

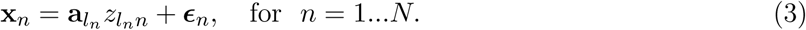

The microstate label of an EEG sample is found as the microstate index, *k*, that minimises the orthogonal squared Euclidean distance,

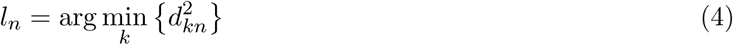

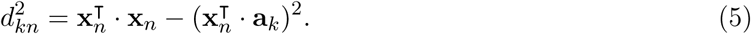

See appendix A for details on this relation as well as an introduction of a new and optimised iteration scheme, for modified K-means. This new iteration scheme upholds the same model assumptions, but preliminary tests show that it is much faster and even slightly better at representing EEG with microstates compared to the original iteration scheme.

The modified K-means method is also accompanied by a scheme for temporal smoothing microstates label sequences to avoid short segments. See section 2.5 for more information on temporal smoothing.

The seminal publication on modified K-means (Pascual-Marqui et al., 1995) contains additional explanation, motivation and pseudo-code.

#### 2.1.3 Topographic Atomize and Agglomerate Hierarchical Clustering

The Topographic Atomize and Agglomerate Hierarchical Clustering (TAAHC) method belongs to the second of two main groups of clustering methods, *hierarchical clustering*. TAAHC was developed for microstate analysis and is a modified version of its precursor, *atomize and agglomerate hierarchical clustering* (AAHC). The (T)AAHC methods differ in significant ways from traditional hierarchical clustering, and even though the only conceptual difference between TAAHC and AAHC is in how they measure the quality of their clusters, their resulting algorithms are significantly different from each other. See section 2.1.4 for a deeper description of these differences.

In TAAHC, the user <u>does not</u> have to pre-set the number of clusters. It starts out with all EEG samples having their own cluster and then one cluster is removed at a time. Each iteration of the algorithm consists of finding the “worst” cluster, and then remove (atomise) it and reassign each of its members to the cluster they are most similar with. This process is then continued until there are only two clusters remaining (or a pre-set minimum number of clusters).

The “worst” cluster is defined as the cluster that has the lowest sum of correlations between its members and prototype (Khanna et al., 2014)^3^:

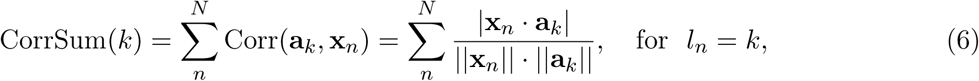

assuming average referenced EEG, i.e. that for each time sample, the channel mean has been subtracted. Note that using the absolute value of the correlation makes TAAHC polarity-invariant^4^. Though the TAAHC is a specialised hierarchical clustering, it can also be seen as a specialised K-means in the way it models the EEG.

By using the correlation to find the worst cluster, the TAAHC is not deterministic (see section 2.1.4) unless an extra step is included in the initialisation of the algorithm. To ensure determinism the clusters are initialised by creating two-sample clusters from the most correlated pairs. In the toolbox this is done by calculating correlations between all sample-pairs, assigning the most correlated pair to the first cluster, and removing the two samples from the sample-pool. This process is then repeated on the remaining sample-pool, until all pairs have been found and the number of clusters has been halved. In the case of an odd number of samples, the leftover sample is assigned its own single-sample cluster.

As a side note on references, Tibshirani and Walther (2005) is often referenced when the (T)AAHC methods are used, but this article actually does not explain how agglomerative hierarchical clustering works. Instead it investigates the relevant question of how to validate the correct number of clusters, when using hierarchical clustering. (T)AAHC is also sometimes erroneously referenced to Pascual-Marqui et al. (1995), which pre-dates the methods and instead introduces the modified K-means algorithm.

For more information on different types of hierarchical clustering we recommend Rokach and Maimon (2005). We suggest using Murray et al. (2008) as a reference for AAHC. Being a new method, the TAAHC unfortunately hasn’t, to the knowledge of the authors, been thoroughly introduced in a published article. Though Khanna et al. (2014) introduces TAAHC, its determinism-ensuring initialisation scheme is not covered.

#### 2.1.4 The difference between AAHC and TAAHC

Each word, from right to left, in the *(Topographic) Atomize and Agglomerate Hierarchical Clustering* methods denotes a specialisation of hierarchical clustering. *Agglomerative hierarchical clustering* is also known as bottom-up hierarchical clustering, since it starts from the bottom with single-member clusters (Rokach and Maimon, 2005).

Conventionally, in agglomerative hierarchical clustering the two most similar clusters are merged in each step, meaning that all of their members now belong to the same merged cluster. However, Murray et al. (2008) argues this may lead to a “snowball effect”, inflating the size of a few clusters. This effect is deemed unwanted as it might interfere with the detection of short periods of stable topography. The AAHC method is specifically designed to counteract this snowball effect.

Instead of merging similar clusters, AAHC finds the “worst” cluster, disbands (atomises) it, and assigns each of its members to the cluster they is most similar with. In AAHC the worst cluster is defined as the one that has the smallest contribution to the quality of the clustering, as measured by the sum of global explained variance^5^ (GEV) of its members (Murray et al., 2008):

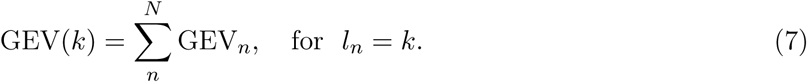

In other words, what makes AAHC different from standard agglomerative clustering is that the members of the removed cluster can join different clusters. This unfortunately also means that you are not able to straightforwardly create a dendrogram, which, though not vital for selecting how many microstates to use, could be used to analyse the structure of the clustered data.

As mentioned earlier, the only difference between TAAHC and AAHC is in how they measure the quality of the clustering. Where the AAHC uses the GEV, TAAHC uses the sum of correlations as seen in (6). This means that TAAHC does not account for the strength of the maps, but only for the similarity of their topographies. Though their definition of cluster quality is the only conceptual difference, it has the effect that the single-member initialisation of TAAHC becomes stochastic, instead of deterministic like AAHC (see section 2.3.1).

TAAHC is stochastic when it initialises using single-sample clusters since all clusters will have a correlation of 1 with their prototypes (because the prototypes are equal to their single member). All clusters are therefore equally bad (or good) and the first “worst” cluster to atomize has to be chosen at random. The AAHC method uses GEV, that weighs the correlation by the global field power (GFP, see section 2.2.1), to find the worst cluster. It is unlikely that two samples have precisely similar GFP, which makes AAHC deterministic. Because of these differences the AAHC has been made available in the toolbox. To ensure determinism, a special initialisation scheme was introduced for TAAHC, as described in section 2.1.3.

Compared to traditional hierarchical clustering it is also an important difference in how the (T)AAHC methods measure similarity (or determine the worst cluster). In standard hierarchical clustering, it is common to use at either the similarity between the cluster prototypes or use the average of all pairwise similarities between cluster members, making the similarity measures independent of the amount of members a cluster has. However, in (T)AAHC the worst cluster is found by <u>summing</u> the correlation or GEV_*n*_ (see (6) and (7)) for each cluster member. This way clusters are “awarded” for having more members, even if they are a bad fit.

So even though AAHC was introduced with the aim of preventing a snowball effect (of large clusters absorbing smaller clusters) it might have inadvertently created a different kind of snowball effect in its method for deciding which cluster to atomise. To our knowledge the effect of these differences have not been investigated in published articles.

#### 2.1.5 Experimental algorithms

We have chosen to collect clustering methods that are not known in the microstate community in a submenu, to avoid cluttering the algorithm selection menu. Currently an unpublished method, *Variational Microstate Analysis*, is implemented in the toolbox, and is selectable under *Experimental algorithms*. This submenu is also intended for future experimental clustering methods for microstate analysis.

### 2.2 Selecting the number of microstates using measures of fit

Though selecting the number of microstates to use is an important choice, it is not a straightforward choice to make. In many situations there is not a single correct answer, but instead many numbers of microstate clusters that are able to explain your data well. One of the issues is deciding how to measure and validate how well your clusters explain your data (see e.g. Tibshirani and Walther, 2005).

Measures of fit are used to estimate how well different microstate segmentations explains (or fit) the EEG, used to estimate the prototypes. In microstate analysis one of the common approaches to decide on the amount of microstates clusters, is to calculate four measures of fit and then make a qualitative decision based on these measures and the quality of the topographical maps of the microstates (e.g. do they look physiologically feasible?).

Since the toolbox makes microstate analysis available in Matlab, it is also possible for users to create their own tests for selecting the amount of microstate prototypes. This would also make it possible to use cross-validation^6^, which is a way control that your segmentation reflects neural activity related to the condition the EEG was recorded under, instead of unrelated recording noise. Briefly, in cross-validation one divides the EEG into a *training set* (e.g. 90% of the EEG), for estimating the microstate prototypes, and a *test set* to test how well the prototypes fit this unrelated data^7^.

Below, we will briefly describe the five measures of fit available in the toolbox and refer to Murray et al. (2008) for further discussion on this topic.

#### 2.2.1 Global explained variance

Global explained variance (GEV) is a measure of how similar each EEG sample is to the microstate prototype it as been assigned to. The higher the GEV the better. It is calculated as;

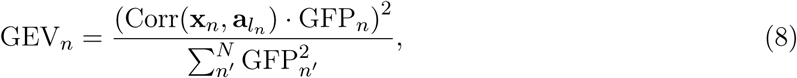

where GFP_*n*_ is the global field power, which is calculated as the standard deviation across all electrodes of the EEG for the *n*’th time sample (Murray et al., 2008). The GEV can be seen as the squared correlation between the EEG sample and its microstate prototype weighted by the EEG sample’s fraction of the total squared GFP:

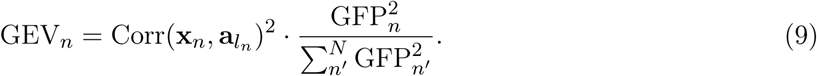

To calculate the GEV for a given cluster, you sum the GEV of each of its members. To use GEV as a measure of fit for a microstate segmentation, you sum the GEV of all samples included in the segmentation.

#### 2.2.2 Cross-validation criterion

The cross-validation criterion (CV) was introduced in Pascual-Marqui et al. (1995). This measure is related to the residual noise, ***∊***, and the goal is therefore to obtain a low value of CV.

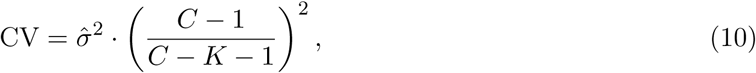

where 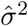m is an estimator of the variance of the residual noise calculated as:

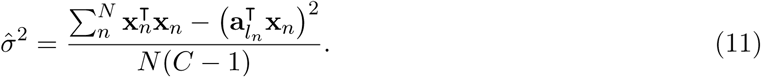

#### 2.2.3 Dispersion

Dispersion (W) is a measure of average distance between members of the same cluster. For a microstate segmentation with K clusters, the dispersion, *W*_*K*_ is calculated as the sum of squares between the members of each microstate cluster:

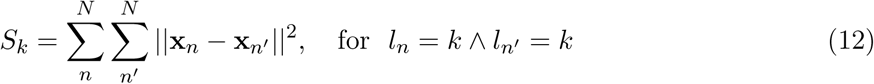

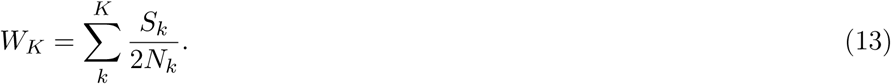

*W*_*K*_ can be considered as an error measure, where the lower the better. It usually decreases monotonically when increasing the number of clusters, since the cluster prototypes can then be closer to its members. It is therefore not a good measure of fit by itself since it will usually encourage using as many clusters as possible.

Note that the sum of squares calculated in (12) is <u>not</u> a polarity invariant measure. Since it takes polarity into account, it might not be a suitable measure of fit for polarity-invariant methods such as modified K-means and (T)AAHC.

#### 2.2.4 Krzanowski-Lai criterion

The Krzanowski-Lai (KL) criterion was introduced in Krzanowski and Lai (1988), as a means of selecting how many clusters to use based on the dispersion measure. High values of KL usually indicates an optimal number of clusters.

Though the *W*_*K*_ usually decreases monotonically there often exists an “elbow”, where the curve flattens. This elbow signifies that there is a drop in the benefit of adding more clusters, and is therefore often chosen as the optimal number of clusters. The KL criterion is a method for detecting such an elbow by looking at the changes in *W*_*K*_ :

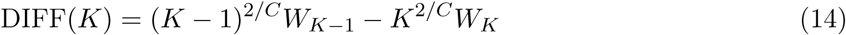

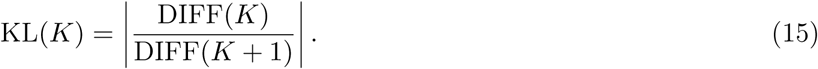

The KL criterion attains large values when an elbow in the *W*_*K*_ curve occurs.

In our toolbox we have added the rule that KL(*K*) is set to zero when *W*_*K*_ − *W*_*K*−1_ is positive, since this indicates that *W*_*K*_ increased instead of decreasing.

Please note, that since the KL is based on *W*_*K*_ it is also not a polarity invariant measure and might not be a suitable measure of fit for polarity-invariant methods such as modified K-means and (T)AAHC. However, since the KL is “just” a method for finding the elbow in the *W*_*K*_ curve, it could be altered to find the elbow in the curve of the measure optimised for by the chosen algorithm. E.g. the modified K-means seeks to optimise the CV criterion, AAHC optimises for GEV and TAAHC optimises for CorrSum(*k*), (6). This feature has not been implemented in the toolbox, but due to the open nature of EEGlab and our an effort to document the code, it should be possible for the user to customise the toolbox.

#### 2.2.5 Normalised Krzanowski-Lai criterion

The normalised Krzanowski-Lai (KL_nrm_) criterion was introduced in Murray et al. (2008) as an adaptation of the original formula, with the intent of computing a normalised curvature of *W*. It is defined as:

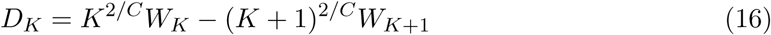

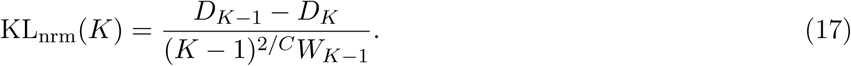

In order to only focus on the convex part of the *W*_*K*_ curve, KL_nrm_ is set to zero when *D*_*K*−1_ *<* 0 or *D*_*K*−1_ *< d*_*K*_.

The main difference between the two KL measures is that the original is a *ratio* of change in the dispersion function, where the normalised version look at the *difference* in the change of dispersion. This means that the two measures are not linearly correlated and that they react differently to dispersion curves. Both measures are therefore made available in the toolbox.

Please note that, like the KL, the KL_nrm_ is also not a polarity invariant measure and might not be a suitable measure of fit for polarity-invariant methods such as modified K-means and (T)AAHC. As a side note on references, Tibshirani and Walther (2005) is often referenced when the KL criterion is used, however this article focuses on using new methods to asses the number of clusters instead of using the KL criterion. We suggest referencing Murray et al. (2008) when using the KL_nrm_ criterion^8^, and the original article, Krzanowski and Lai (1988), when using the KL criterion.

### 2.3 Settings for microstate clustering

Most clustering algorithms has some settings that influence how they perform in segmenting data. In this section we will explain what some of the important settings of clustering algorithms do, and why they are relevant.

#### 2.3.1 Multiple restarts of algorithms

In the introduction of this methods section it was mentioned that some algorithms gives a different result every time they are run, even with the same settings. To explain the reason for this it is necessary to make a distinction between *deterministic* and *stochastic* algorithms.

As the name implies, a deterministic algorithm is an algorithm that always give the same result, when its settings and the data stay the same. In other words there is no randomness in the algorithm. If there is randomness somewhere in the algorithm, it is stochastic. This randomness is often introduced in how the algorithm is initialised.

When using K-means or modified K-means for microstate analysis, the cluster prototypes are often initialised by selecting EEG samples. There is usually not one true way of selecting which EEG samples to use for prototypes, so a common approach is to select the EEG samples at random, making the algorithm stochastic.

Though it would be tempting to always select the EEG samples in the same manner to ensure determinism, it runs the risk of obtaining an inferior result. By restarting the stochastic algorithm multiple times, you are able to test multiple segmentations on the same dataset and select the best restart based on e.g. GEV. The downside is that you are not guaranteed to obtain the same segmentation every time the algorithm is run.

How many restarts to use is a trade-of between computation time, and how likely you are to converge on the same optimal solution. In the toolbox 10 restarts are used by default.

#### 2.3.2 Stopping criteria: Convergence and maximum iterations

Some algorithms, like K-means, work by repeating multiple steps in iterations, which means that it will keep iterating until some stopping criteria is met. One of these criteria is the convergence threshold, which stops the algorithm when the relative change in error between iterations drops below the threshold. By default this is set to 10^−6^ in the toolbox.

Again there is a trade-of between computation time and precision of the segmentation, for which reason it is also possible to set a maximum number of iterations before stopping. By default the toolbox has a maximum number of iterations set to 1000.

### 2.4 Backfitting microstates to new EEG

When you have obtained microstate prototypes from your chosen clustering method, you might wish to see how well some other recordings fit with these prototypes. This is relevant, e.g., when working with spontaneous recordings, where you can’t reduce the noise levels through averaging trials. Instead you only give the clustering algorithm time samples that peaked in the GFP time curve, as they are assumed to have the “cleanest” representations of their microstate. After obtaining the prototypes it is then relevant to see which microstate all prototypes likely belong to.

Backfitting is assigning a microstate labels to EEG samples based on which microstate prototype they are most topographically similar with. This similarity can be measured using global map dissimilarity (GMD).

The GMD, which is also known as DISS (Murray et al., 2008), is a distance measure that is invariant to the strength of the signal and instead only looks at how similar the topographical maps look. For two EEG samples, **x**_*n*_ and **x**_*n*_′, GMD is calculated as:

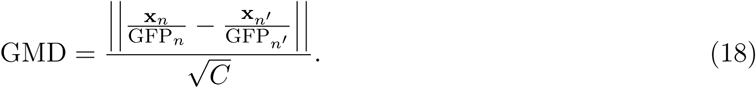

By normalising with GFP, two EEG samples that belong to the same microstate, but have different strength, will achieve a low GMD distance.^9^

### 2.5 Temporal smoothing

EEG recordings are known to contain a lot of unwanted noise, especially in spontaneous recordings where the noise can’t be averaged out like with ERPs. This noise can contribute to short (i.e. few or single-sample) microstate segments appearing after clustering or backfitting. One way to address this is to use temporal smoothing, where an EEG sample is not only assigned to a microstate class based on topographical similarity with the prototype, but also based on the microstate labels of samples prior and following the EEG sample.

The most commonly used clustering algorithms in microstate analysis do not take the temporal order of the EEG samples into account. Therefore the temporal smoothing will be done as a processing step after the microstate segmentation has been run.

Currently there are two variations of post-segmentation smoothing implemented in the toolbox; *windowed smoothing* and *small maps rejection*.

#### 2.5.1 Windowed smoothing

This variation of temporal smoothing of microstate segments was introduced in Pascual-Marqui et al. (1995).

The microstate segments are smoothed by updating the microstate labels using a distance measure, that is a trade-off between optimising the best fit to data and temporal smoothness of the labels.

This distance measure is obtained by adding a bonus term for temporal smoothness to the distance measure expressed in (5);

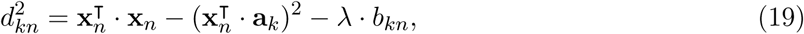

where *b*_*kn*_ is equal to the amount of samples that has the microstate label *k* in a window surrounding the *n*’th sample, and *λ* denotes how strongly to weigh smoothness.

This method of smoothing needs the user to set two parameters; *λ* and the width of the window used. Pascual-Marqui et al. (1995) suggets to set *λ* to 5, and the window width to three samples (on each side of the current sample), however, the optimal settings might vary between datasets, e.g., due to varying sampling frequency or recording conditions. To find the optimal smoothing settings it would therefore be necessary to use cross-validation.

#### 2.5.2 Small segment rejection

For this toolbox, we introduce a method which sets a minimum duration microstate segments are allowed to last. The algorithm repeatedly scans through the microstate segments and changes the label of time frames in such small segments to the next most likely microstate class, as measured by GMD. This is done until no microstate segment is smaller than the set threshold.

### 2.6 EEG microstate statistics

Parsing the EEG data into microstates (i.e. labelling of the EEG data with respect to the best fitting microstates classes) offers a rich set of statistical parameters with potential neurophysiological relevance. These statistical parameters can be divided into parameters about the activation strength, the spatial configuration and the temporal attributes of microstates. The strength of the average global activation during a given microstate *k* is defined by its *average* GFP of all EEG samples assigned to microstate *k* :

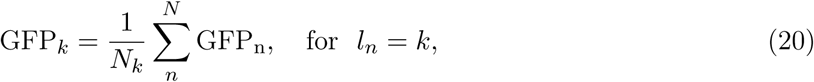

where *N*_*k*_ is the number of samples assigned to cluster *k*. See section 2.2.1 for more information regarding GFP.

The *average* GEV (see equation 8 and 9) and the *average spatial correlation* (between microstate prototype maps and their assigned EEG samples) reflect to what extent the microstate protoypes can explain the data. Where GEV looks at both the strength of the EEG and the spatial fit, the average spatial correlation only looks at the spatial fit.

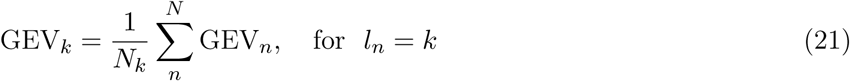

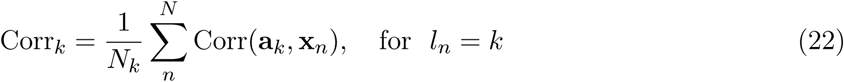

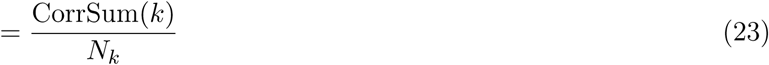

Note that in the toolbox the correlation is polarity-invariant, i.e. non-negative.

The basic temporal dynamics of microstates are described by Occurrence(*k*), Duration(*k*), and Coverage(*k*). In the toolbox these are calculated using run-length encoding. Occurrence(*k*) reflects the average number of times per second a microstate is dominant, the Duration(*k*) is defined as the average duration of a given microstate (in milliseconds), and the Coverage(*k*) reflects the fraction of time a given microstate is active.

In addition to these basic statistical parameters, the *transition probabilities* between microstates can be derived to quantify how frequently microstates of a certain class are followed by microstates of other classes. There are (at least) two variants of this statistic, one where the transition probabilities are not corrected for different number of occurrences of microstates (i.e. base rates of microstates) and one which adjusts for the base rate probabilities. Note that only the former version is currently implemented in the toolbox.

## 3 Guide to the Toolbox

In this part, we provide a practical guide how to conduct EEG microstate analyses using the EEGlab toolbox. Figure 1 provides an overview of the specific processing steps pertinent to the analysis of spontaneous EEG data and for event-related potential (ERP) data with the corresponding `Matlab functions` and typical arguments that have to be passed to the functions. In the subsequent sections we will provide a more thorough tutorial that shows how to analyse a small dataset of spontaneous EEG data (see 3.1). We would like to emphasise that our choices regarding algorithms, settings, and even the whole work-flow itself are based on our experience and not on empirical evidence, which is, to the best of our knowledge, lacking so far.

**Figure 1:**
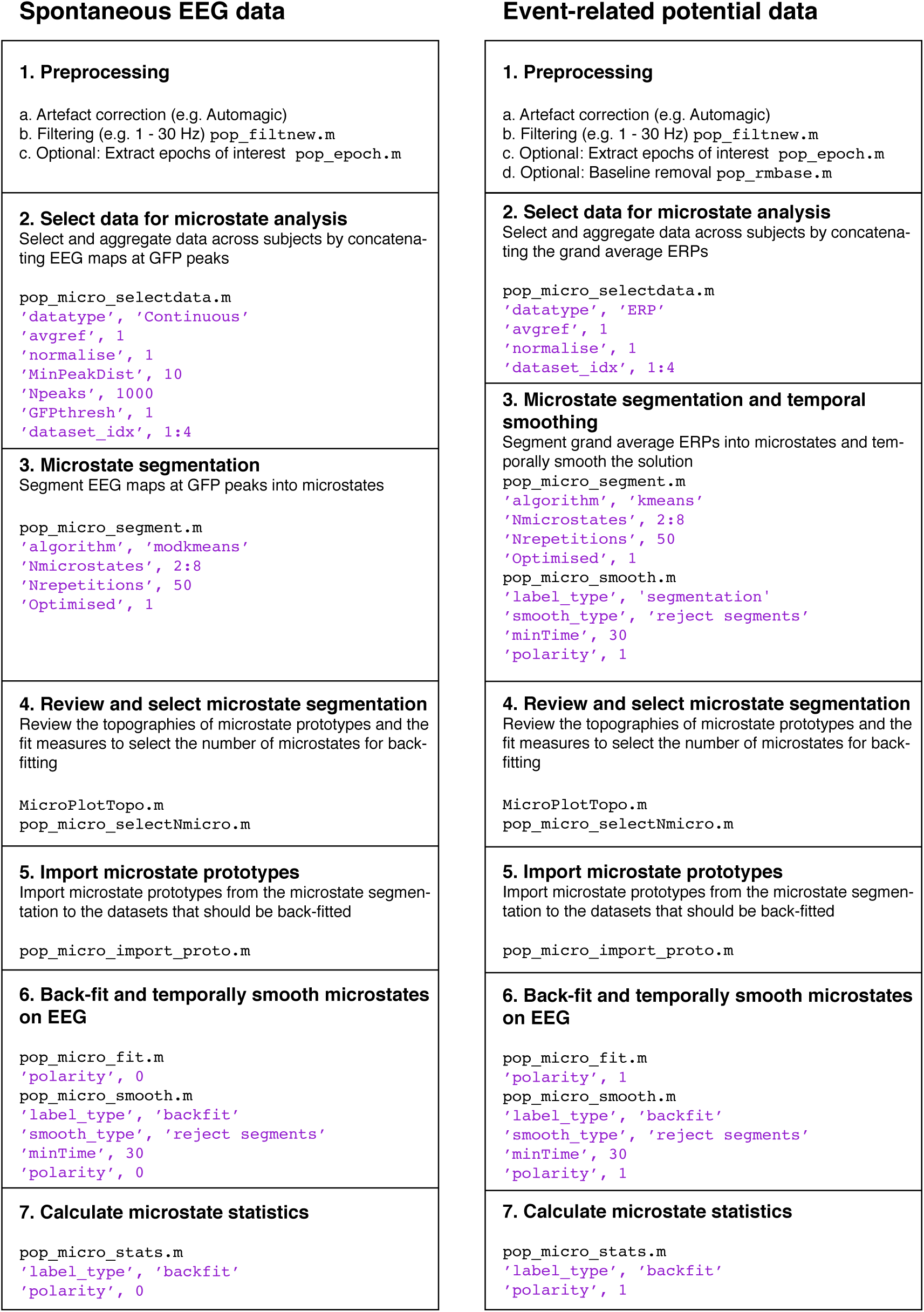
Schematic overview of a workflow for spontaneous EEG and ERP data.

### 3.1 Tutorial: Microstate Analysis on spontaneous EEG data

This tutorial will demonstrate how to use the graphical user interface (GUI) of the Microstate Toolbox for EEGLAB to perform a microstate analysis on a small set of spontaneous EEG data with four subjects. The data used for this tutorial can be downloaded here: https://archive.compute.dtu.dk/files/public/users/atpo/Microstate(∼ 350 Mb). The archive also contains the newest version of the toolbox and this guide. With this dataset, it should be possible to reproduce the actions discussed in the tutorial and to obtain similar results as shown in the figures and outputs.

To be able to use the Microstates Toolbox presented in this tutorial, Matlab and EEGLAB (Delorme and Makeig, 2004) needs to be installed and the tutorial data should be stored in an accessible folder. To install the toolbox, simply unzip the toolbox into your MATLAB/eeglab/plugins folder. The toolbox is tested with Matlab version 2015b or later and EEGLAB version 14.0.0 or later.

#### 3.1.1 Description of the tutorial dataset and completed preprocessing steps

The tutorial data consists of EEG from 4 subjects (with the original IDs: A00062279001, A00062578001, A00062842001, A00062219001) of the age group 25-44 from the data sharing paper by Langer et al. (2017)^10^. The datasets have been preprocessed using the Automagic^11^ artefact correction (with the standard settings) and have been filtered with a highpass filter of 1 Hz and a lowpass filter of 30 Hz using the standard settings of pop_firfiltnew() from EEGLAB 14.0.0b.

#### 3.1.2 Automatising analysis steps for multiple datasets

After finishing this tutorial, the reader may imagine that conducting the analysis steps using the GUI for larger datasets may become a tedious and an error-prone endeavour. A more efficient and reproducible way of conducting the same analyses can be accomplished using a MATLAB script that calls the functions that perform the specific processing steps for all datasets as a batch process. Such scripts can be easily compiled using EEGLAB’s history command `eegh`^12^. Furthermore, a script that performs the exact analysis steps of this tutorial is provided in appendix B.

### 3.2 Outline

To recap, the goal of a microstate analysis is to parse EEG maps into microstate prototypes and re-express the spatio-temporal characteristics of the EEG time series by means of the microstate prototypes. Thus, the first step (after loading the data) is to derive a set of microstate prototypes by clustering the EEG data into as few as possible microstate prototypes that account for as much as possible variance in the EEG data. For the microstate analysis of spontaneous EEG, such as resting state data (as opposed to time-locked EEG data), the amount of data is typically reduced by only using the EEG maps, where the GFP is peaking. This reduces the number of EEG maps that enter the analysis (and hence the processing time) and discards maps of low signal-to-noise ratio. The selected EEG at GFP peaks enters a clustering algorithm that groups it into a small set of classes based on topographic similarity and calculates for each microstate class a topographical prototype, the so called microstate prototype.^13^ After obtaining the microstate prototypes, the topography of each EEG sample is labelled as belonging to one of these classes, and the EEG signal is re-expressed as a sequence of microstate classes. Finally, statistics about the sequence of microstate classes, such as their frequency of occurrence or average duration can be calculated. These statistics are typically submitted to further statistical analyses such as group comparisons.

### 3.3 Data selection and aggregation

#### 3.3.1 Loading datasets in EEGLAB

Start EEGLAB and load each of the four datasets with `Load existing dataset`^14^. This step has to be repeated for each dataset. All loaded datasets can be viewed and selected from the `Datasets` menu (see Figure 2).

**Figure 2:**
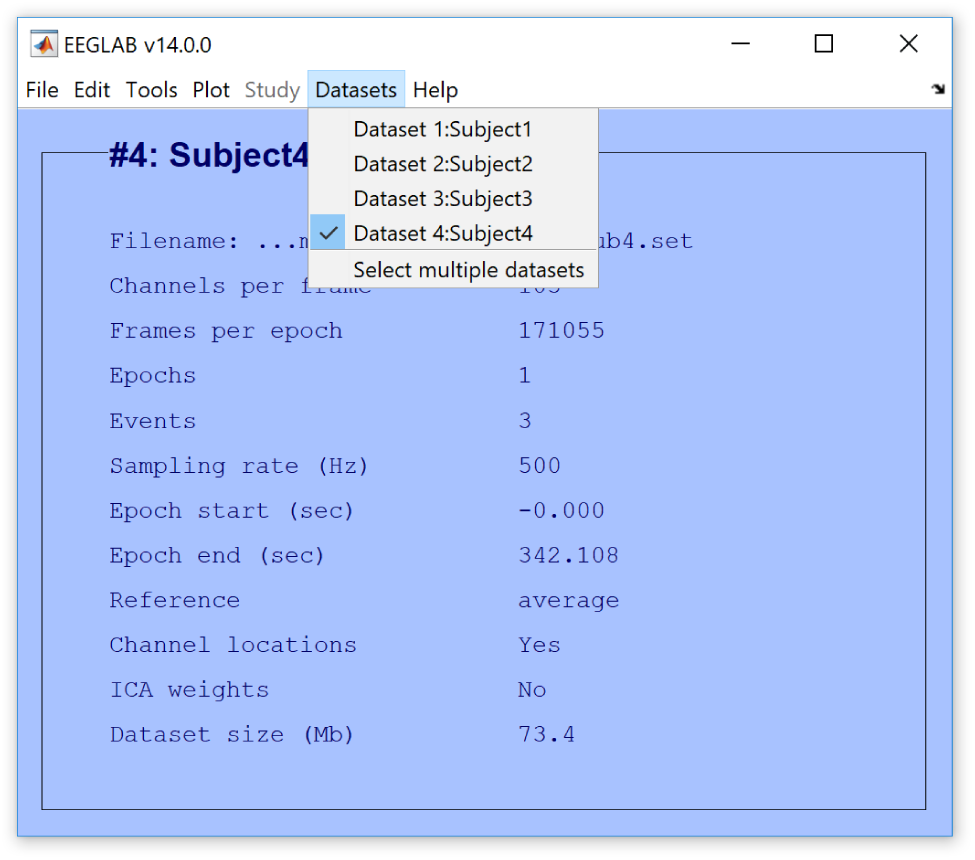
Datasets loaded into EEGLAB.

#### 3.3.2 Select data for microstate segmentation

Before starting the microstate segmentation, it is necessary to specify the data of which the microstates will be derived from. Since, we are typically interested in obtaining microstate prototypes that explain variance in datasets consisting of more than one subject, the toolbox provides functions to aggregate EEG from multiple datasets into one dataset. This dataset is then used to derive microstate prototypes from.

To select data, open the menu `Tools/Microstates/Select data for microstate analysis`. This will open a window (Figure 3a), which allows you to select the following options (with suggested settings):

- *Data type*: By selecting *Spontaneous - GFP peaks*, EEG maps that represent peaks in the GFP time curve will be used for segmentation. By selecting *ERP*, EEG from all time points will be used. If the dataset contains multiple epochs/trials, these will be averaged into one average epoch (i.e. a grand average ERP). Since the data used for this tutorial stems from spontaneous EEG, we set the data type to *Spontaneous*.
- *Select and aggregate data from other datasets*: This option allows to perform the microstate segmentation on EEG from multiple datasets. In the case of *Spontaneous* data, for each subject, a selection of defined number of EEG maps at GFP peaks is randomly extracted and concatenated to one new dataset^15^. This option will open a new window, where the datasets that are already loaded into EEGLAB are listed and can be selected. Note, if this option is not selected, the segmentation is only performed on the currently active dataset.
- *Calculate average reference*: We advise to reference the data to average reference. Michel et al. (2009) discuss why it is a good idea to use the average reference.
- *Normalise dataset(s)*: This option will normalise each dataset with by the average channel standard deviation within each dataset. This makes the amplitude of the EEG comparable between recordings, which might help weighing each dataset equally in the microstate clustering. However, if the data contains high amplitude artefacts the normalisation might skew the weight given to each dataset. For this walk-through we will not select this option.
- Settings for the extraction of GFP peak maps - These are only relevant and active when selecting *spontaneous* data:

– *Minimum peak distance (ms)*: Defines what the minimal distance between GFP peaks can be. This is to ensure only using distinct peaks. We set the minimum peak distance to 10 ms.

**Figure 3:**
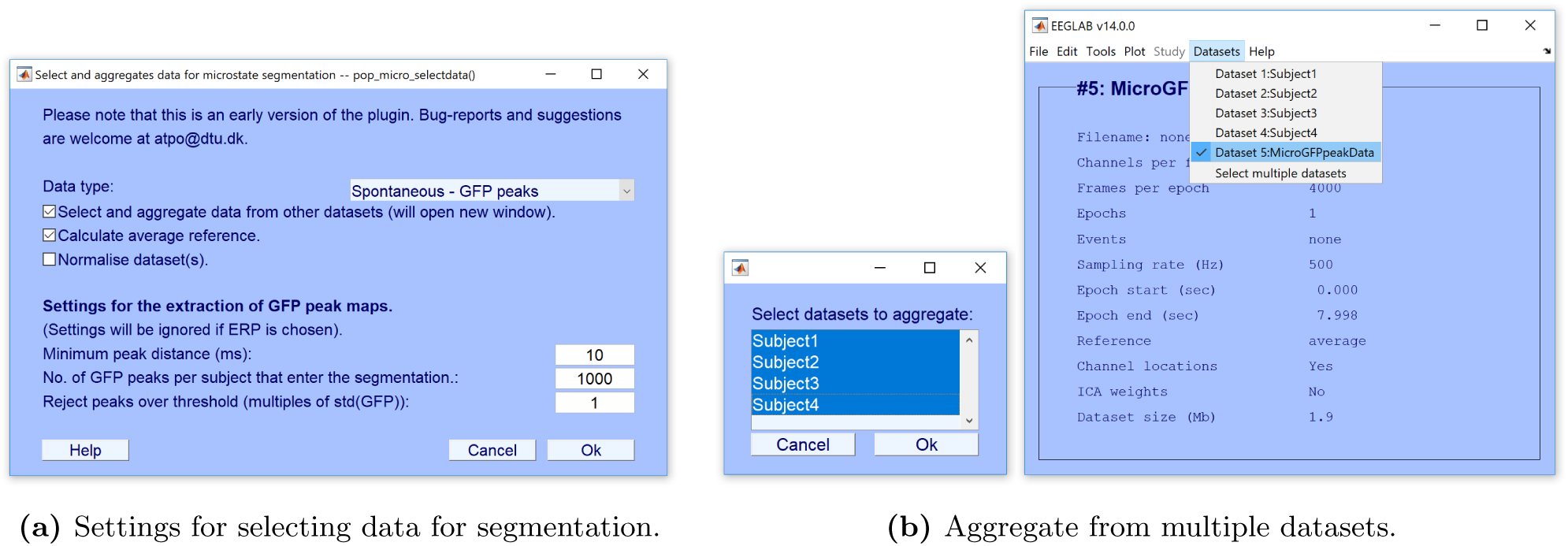
Select your settings for the data to be segmented. If you are aggregating from multiple datasets, another window will prompt you to select which datasets to aggregate from, and a new dataset will be created.
– *No. of GFP peaks per subject that enter the segmentation*: Indicates how many GFP peaks are maximally extracted per dataset. We choose to extract EEG samples from 1000 randomly selected GFP peaks for each subject.
– *Reject peaks over threshold* : Discards maps with a GFP that exceeds X times the standard deviation of the GFPs of all maps. This option exists because maps with extreme GFPs often include artefacts of non-neural origin with high amplitudes. Here we will reject maps with GFP that exceeds 1 times the standard deviation of GFPs. Note that this rather low threshold prevents from an unwanted influence of outliers that can occur when analysing data from only four subjects. If there are few GFP peaks available per subject, it may be reasonable to select higher values for this setting.

Confirm the settings by clicking *Ok.* A new window *Select datasets to aggregate* (see Figure 3b) appears to select the datasets to include in the analysis. Use the Shift or Ctrl keys to select multiple datasets and confirm your selection by clicking *Ok.* A new dataset will be created called *MicroGFPpeakData* (which is now listed under `Datasets`) that includes the EEG extracted and aggregated based on the GFP peaks.

Please note, that we are going to estimate microstate prototypes based on GFP peaks selected at random, and there will therefore likely be slight differences in the resulting microstates compared to the ones illustrated on figures 5b and 9b. We experienced converging results when using 1000 peaks per subject for this specific dataset. However, note that for other datasets it may be necessary to take more GFP peaks.

**Figure 5:**
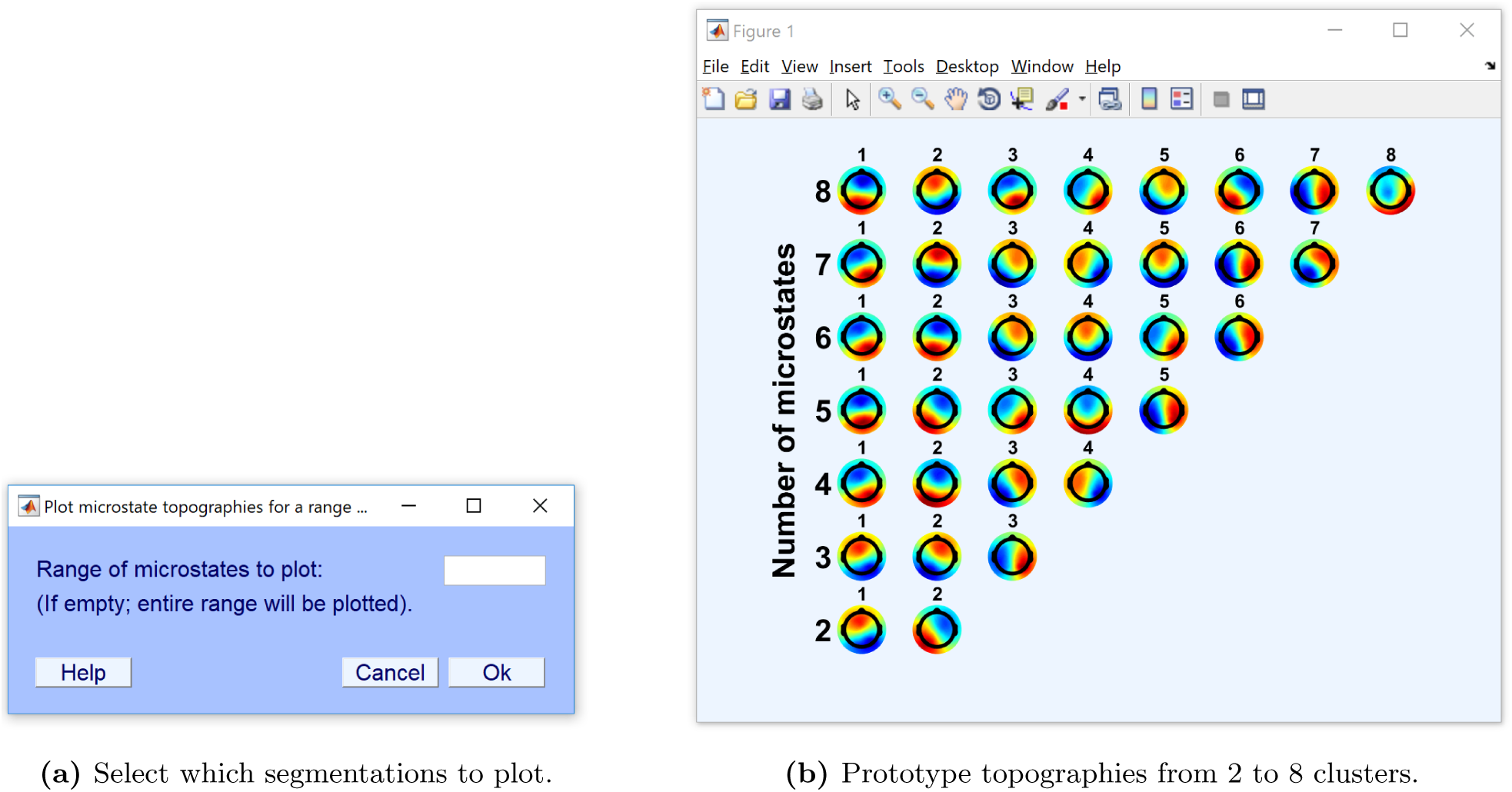
A qualitative assessment of the prototype topographies can be used to decide how many clusters to use. Note that since modfied K-means is polarity-invariant the prototypes for different segmentations might be inverted as is the case for prototype number 1 for 2 and 3 clusters.

### 3.4 Microstate segmentation

In this step, the selected EEG (in our case aggregated GFP peaks from four subjects found in *Datasets: MicroGFPpeakData*) is segmented into a predefined number of microstate prototypes with the goal of maximising the similarity between the EEG samples and the prototypes of the microstates they are assigned to. How this similarity is measured is dependent on the chosen clustering algorithm (see section 2.1).

Make sure that the dataset *MicroGFPpeakData* is the currently active dataset (indicated by a tick in the menu `Datasets`) before opening the menu `Tools/Microstate analysis/Segment into microstates` to select (Figure 4a).

**Figure 4:**
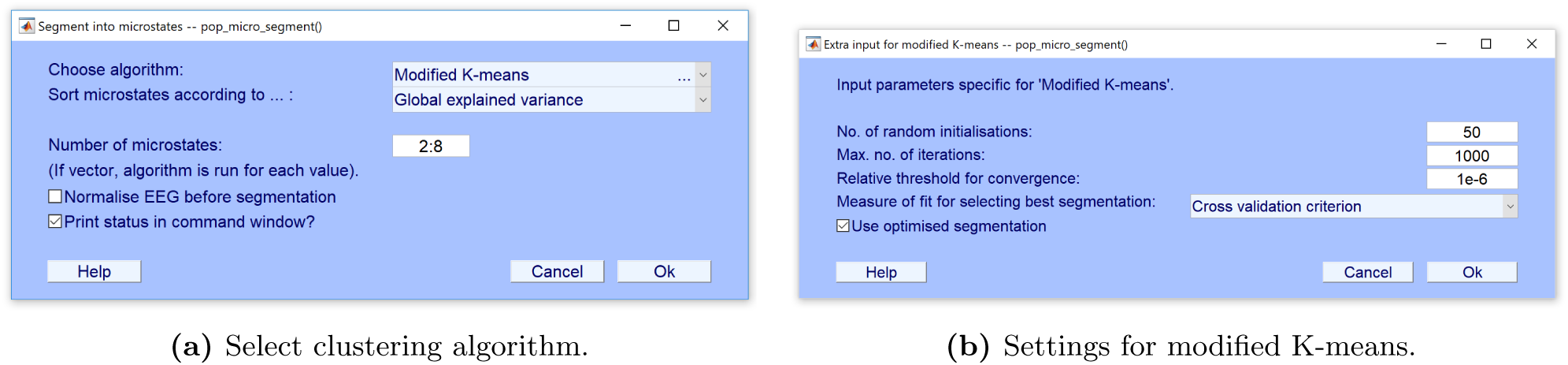
Settings for microstate segmentation. After selecting a clustering algorithm, another window will appear with settings specific to the selected algorithm.

As we are working with spontaneous EEG data, we will choose the *modified K-means* algorithm^16^, since it ignores the polarity of the EEG topography (Pascual-Marqui et al., 1995; Michel et al., 2009; Wackermann et al., 1993; Lehmann, 1971).

By default the microstate prototypes are sorted by decreasing *Global explained variance*. The sorting is done after the clustering and does not affect the segmentation. We set the sorting to its default setting and set the *Number of microstates* to cluster the EEG data into from 2 to 8 microstate prototypes (which seems reasonable considering that a majority of studies find four microstate prototypes to be most adequate to describe resting EEG data).

It is possible to *Normalise EEG before segmentation* by the average channel standard deviation, as was the case in `Select data for microstate analysis`. However, here the normalisation will be done across the entire dataset and only for the segmentation, i.e. the stored data will not be changed. This could be relevant, if one wanted to keep the dynamic ranges between datasets by not normalising during data aggregation, but still wanted to help the clustering methods that sometimes assume unit standard deviation. For the same reasons as earlier, we will leave the option unchecked for this walk-through.

After confirming the settings with *Ok* an additional window will open (Figure 4b), in which we can set parameters for the modified K-means algorithm. As mentioned in section 2.3 the first three settings reflect a trade-off of reproducibility versus computation time.

In general, it is advisable to set as high an amount of *No. of random initializations* as computationally feasible to obtain a replicable result. This can be tested by repeatedly running the segmentation and comparing the topographies of the resulting microstate prototypes. If the algorithm reaches the pre-set *Maximum number of iterations* for all restarts (as can be observed in the output on the command window in Matlab), it is recommended to increase this value. The *Measure of fit for selecting best segmentation* defines how to decide which of the *N* restarts obtained the best clustering. As modified K-means seeks to optimise the clustering based on the CV criterion, we will also set this as the measure of fit.

We *Use optimised segmentation* to employ the optimised iteration scheme for modified K-means that is introduced in appendix A. We use this new iteration scheme, since our preliminary tests indicate that it is much faster and is slightly better at representing the EEG as microstates. The excat computational speedup depends on the dimensionality of the segmented data, but for the present data the optimised iteration scheme results in a computational speed-up by a factor of ∼5. We will use this speed-up to increase the *Maximum number of iterations* to 50 and leave the other two settings to their default values.

### 3.5 Review and select microstate segmentation

After having clustered the EEG with multiple numbers of microstate clusters it is necessary to select which number of clusters to use for further analysis. In microstate analysis, this is often decided based on a qualitative evaluation of the prototype topographies and measures of fit. In some study that analyse spontaneous EEG, while participants are at rest, 4 clusters are chosen, based on previous results, summarized in (Michel and Koenig, 2017).

#### 3.5.1 Plot microstate prototype topographies

To review the result of the microstate segmentations for different numbers of microstates, we look at the quality of the microstate prototypes topographies by selecting `Plot microstate prototype topographies`. This opens a figure as seen on Figure 5b that shows the microstate prototype topographies of each microstate segmentation, sorted by the measure selected in `Segment into microstates` from left to right. The qualitative assessment of the microstate prototypes can, e.g. be based on if the same microstate segmentation contains prototypes with very similar topography or whether the topographies seem physiologically feasible.

#### 3.5.2 Select active number of microstates

In addition to selecting the number of microstates based on the topographies, we should review the quality of the different microstate segmentations with respect to the goodness of fit. There are various measures for this as explained in section 2.2. Since these measures are calculated in different ways, segmentations that score well with one measure do not necessarily score just as well with other measures. Importantly, the fit measures W, KL and KLnorm are not polarity-invariant, as it is assumed in the segmentation of spontaneous EEG data. Therefore, we should only use the GEV and the CV criterion to review the goodness of fit of our microstate segmentations. To do this, open the menu `Tools/Microstate Analysis/Select active number of microstates` (Figure 6a) and deselect W, KL and KLnorm and only select GEV and CV and click *Ok*.

**Figure 6:**
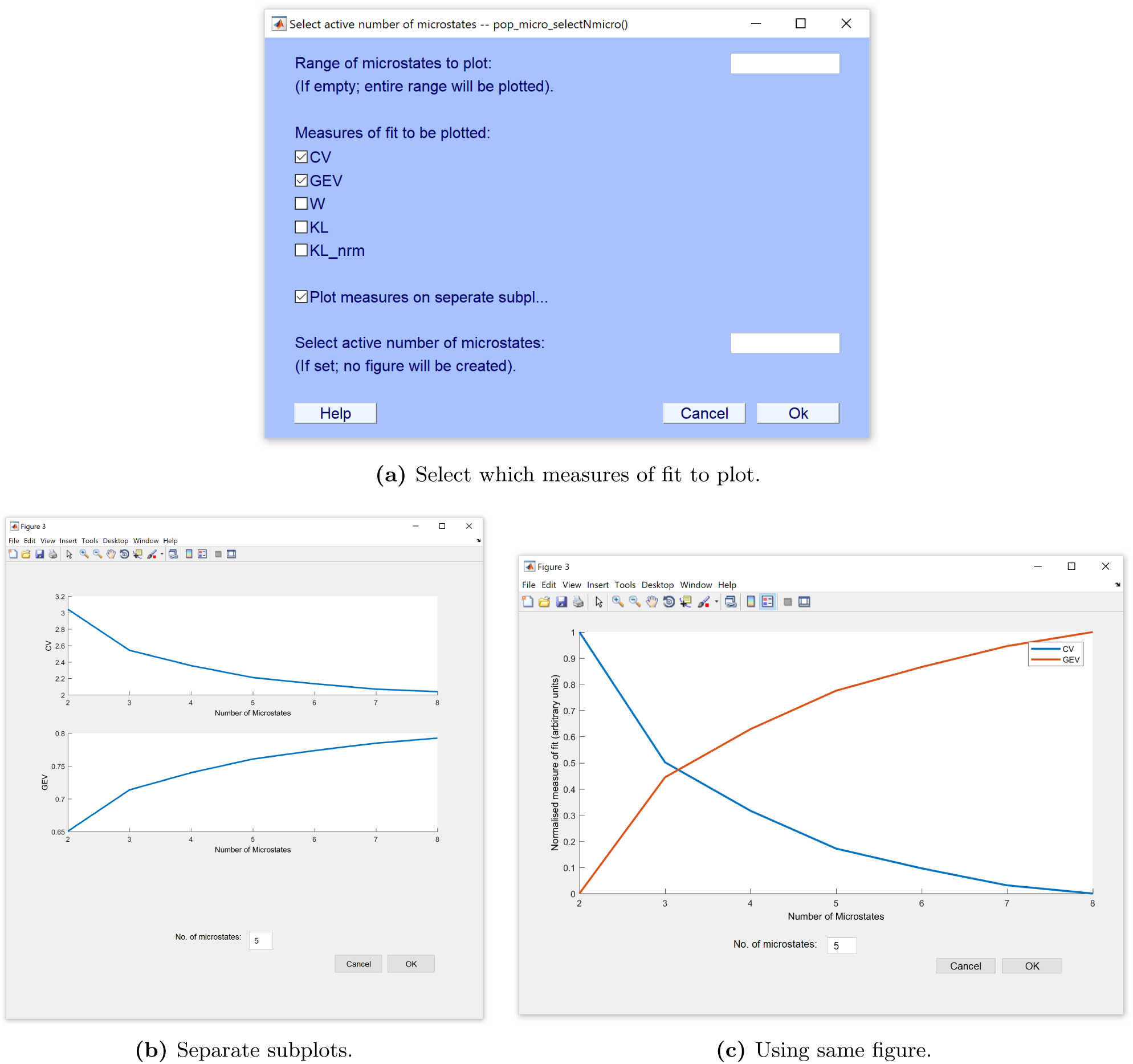
Select active number of microstates based on measure of fit. These can either be plotted on individual subplots or in the same figure. If plotted together, the plotted measures of fit are normalised to lie in the range [0; 1]. When clicking *Ok*, the segmentation from with the *No. of microstates* will become the active segmentation.

A new figure shows the measures of fit plotted for the different microstate segmentations. This is either plotted on subplots (Figure 6b) or normalised together (Figure 6c), depending whether we have previously selected *Plot measures on separate subplot*. If plotted together, each measure is normalised independently to lie between 0 and 1.

The decision for the right number of clusters obviously reflects a trade-off between the goodness of fit and the complexity a high number of microstates brings to the segmentation. The GEV criterion theoretically (and most of the time effectively) becomes monotonically larger, when increasing the number of clusters, and it is therefore a question of stopping when adding another cluster does not bring a significant benefit.

Note that it can happen that the GEV does not monotonically increase. This indicates that the segmentation has not arrived at the best clustering solution, which might be overcome by increasing the number of restarts and maximal number of iterations and decreasing the threshold for convergence to ensure the best solution is obtained for each number of microstates. while the GEV does not account for complexity (i.e. degrees of freedom), the CV criterion does. However practically, the CV criterion, pointing to the best clustering solution at its smallest value, often reaches such a minimum only with large numbers of clusters, which are difficult to interpret.

In our case an argument could be made for choosing between four and six clusters. We select using five clusters since it has a good balance between goodness of fit and the number of clusters, as well as it obtains feasible topographies.

### 3.6 Back-fit microstates on EEG

Now that we have selected the number of microstate prototypes, we are ready to fit the microstate prototypes back to all of the EEG (and not only the GFP-peak data we selected for segmentation). In this processing step, we essentially label the EEG data with the class of the microstate prototype that is most similar. To do this we need to import the microstate prototypes to the dataset to which the prototypes are backfitted to.

We start by changing the active dataset in `Datasets` to the first dataset (Subject1). We want to import the microstate prototype topographies we just obtained, which is stored in the dataset *MicroGFPpeakData*. We therefore select `Import microstate prototypes from other dataset` and then *MicroGFPpeakData* from the list of datasets (see Figure 7). This will add the field `microstates. prototypes` to the `EEG` structure in the Matlab workspace.

**Figure 7:**
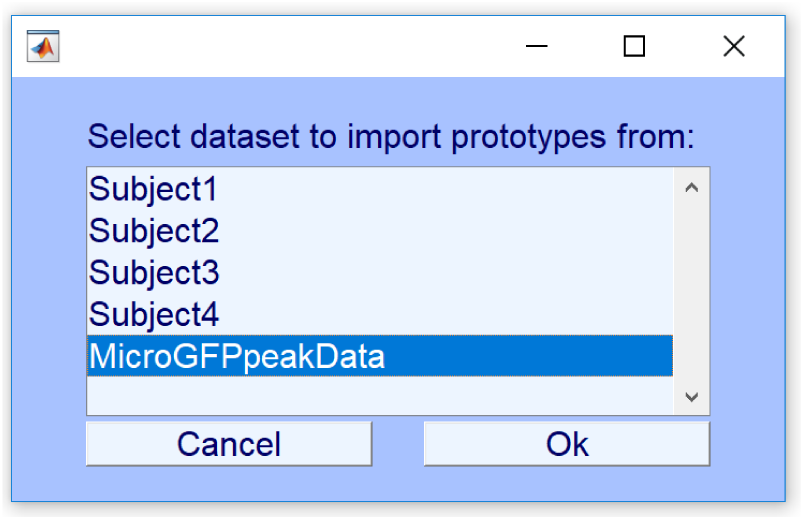
Import microstate prototypes from the *MicroGFPpeakData* dataset into the current dataset.

As a last step in the back-fitting, we select `Backfit microstates on EEG` to open a window, in which we can choose whether we want to ignore polarity when back-fitting. As we are analysing spontaneous EEG, we want to ignore polarity. So leave this box unchecked and press *Ok*.

### 3.7 emporally smooth microstate labels

In the back-fitting procedure, each EEG sample is assigned to the class of the microstate prototype it is most similar with. As spontaneous EEG includes noise, it will happen often that consecutive time frames are labelled differently by chance. In addition, short periods of unstable EEG topographies occur typically during the change of polarity, even in the same microstate. To reduce these spurious influences, microstate labels can be temporally smoothed after the back-fitting. To do this we open the menu `Temporally smooth microstates labels` (See Figure 8a).

**Figure 8:**
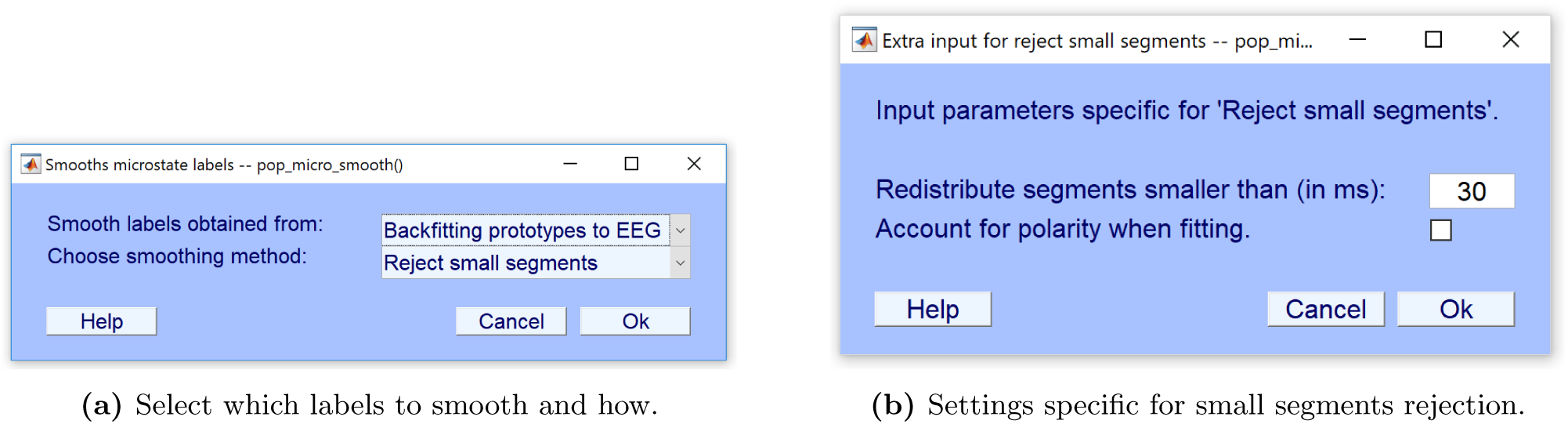
Temporal smoothing microstate labels. The microstate labels can either stem from prototypes backfitted to EEG, or from the EEG used in the microstate clustering.

In *Smooth labels obtained from:* select *Backfitting prototypes to EEG*. Here we select the smoothing method *Reject small segments*.^17^ Click *Ok*.

A window opens (Figure 8b), where we can indicate how small microstate segments that we will tolerate before the EEG samples are redistributed to other microstate clusters. We use the default settings and click *Ok*.

### 3.8 Illustrating microstate segmentation

One might wish to visualise the microstate segmentation. This works especially well for ERP analysis, where the entire epoch can fit in a single figure. Since we use spontaneous data for this guide we will instead visualise 1.5 s from subject 1. Figure 9 shows the GFP of the active microstates in the time range between 4,200 and 5,700 ms of the entire resting EEG dataset for subject 1.

**Figure 9:**
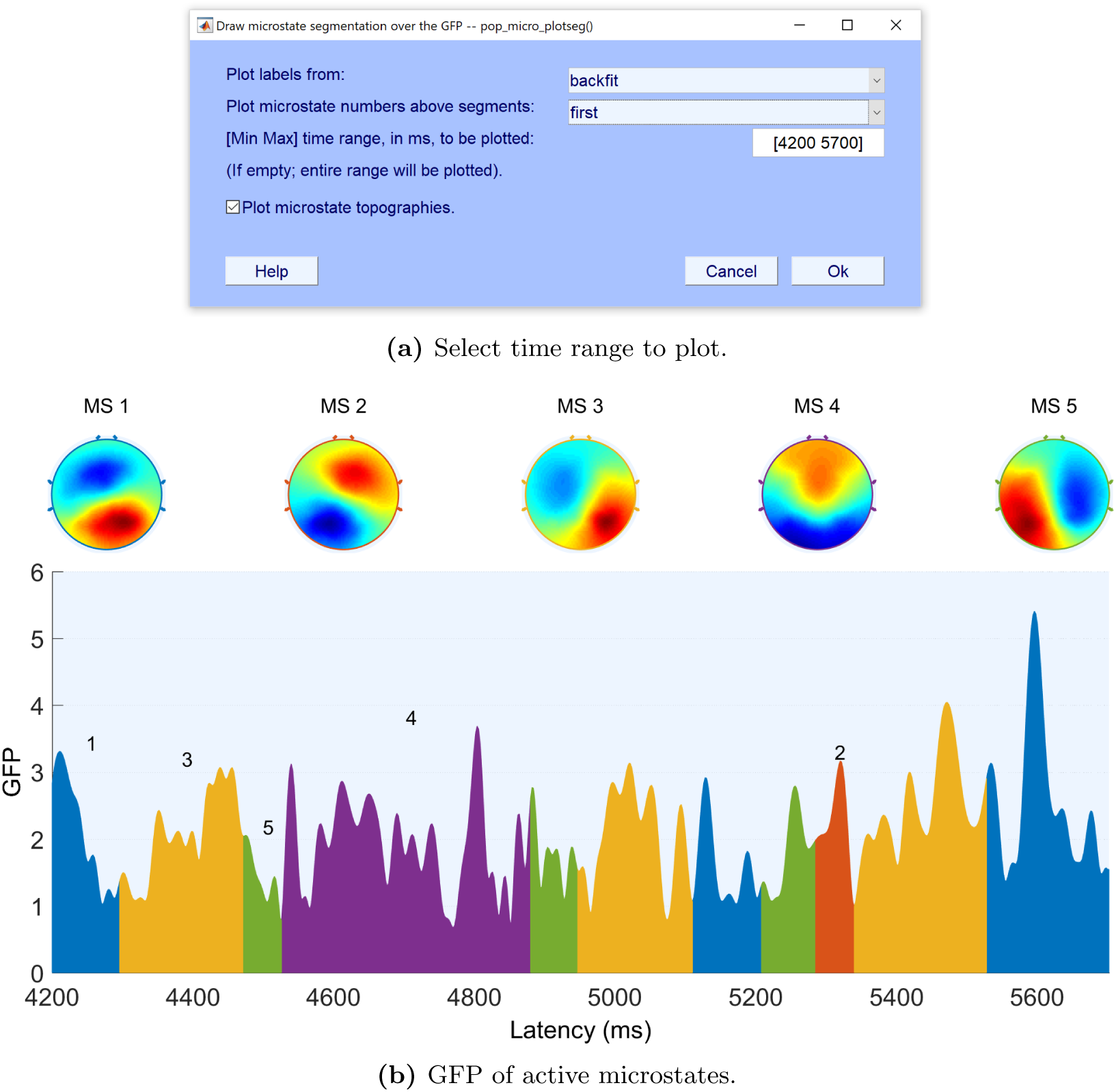
Illustrative figure of the GFP of active microstates for 1,500 ms, from 4,200 ms to 5,700 ms, of the EEG from the first subject.

### 3.9 Calculate microstate statistics

Once we have back-fitted to all of the EEG and temporally smoothed the microstate labels, we can calculate the statistical properties of the microstates (see section 2.6), including the average GFP in all time frames of the same microstate class, the occurrence of each microstate class per second, the average duration, the percentage coverage of each microstate class, the Global Explained Variance and spatial correlation (i.e. on average how much variance in the EEG data is explained by the best fitting microstate prototype) and the transition probabilities between microstate classes. To run these calculations click on `Calculate microstate statistics`. This will open a window (Figure 10), in which you select *Use labels obtained from Backfitting prototype* to EEG and leave the entry for *Time window of analysis* empty, so it uses all time frames. Leave the option *Account for polarity when fitting* unchecked, as we have not accounted for polarity previously in the analysis.

**Figure 10:**
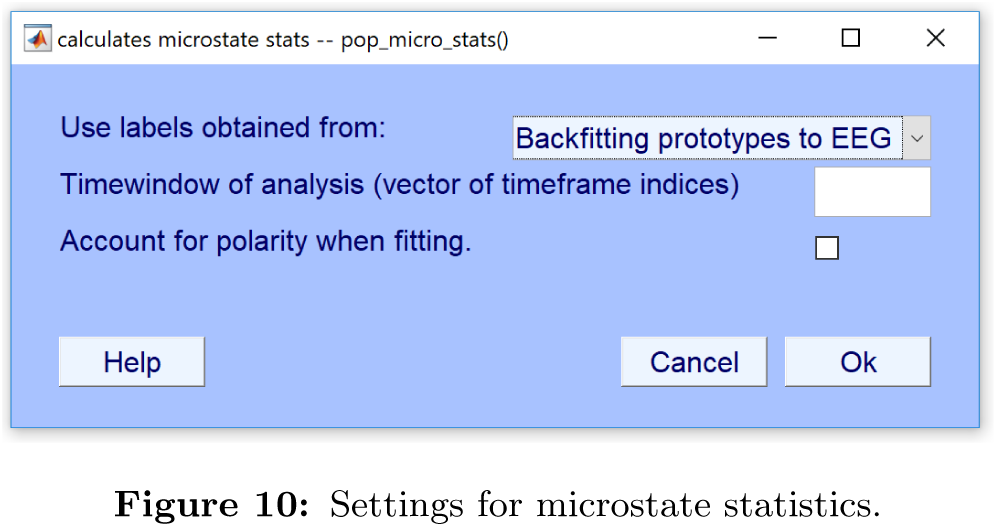
Settings for microstate statistics.

The results of the microstate statistics are stored in the `microstate.stats` field of the `EEG` structure in the Matlab workspace. It is then possible for the user to submit these statistics to further custom analyses.

Repeat all steps in sections 3.6, 3.7 and 3.9 for each dataset. See section 3.1.2 on how to automatise these steps to avoid going through the steps in the GUI for each dataset.

## 4 Concluding remarks

The aim of both the toolbox and guide has been to make it easier to adapt to microstate analysis for researchers new to the field. Both by bringing transparency to the employed methods, and by supplying an open source toolbox in Matlab.

It is our impression that researchers that are interested in microstate analysis hesitate to adopt to the field, due to a lack of understanding of the underlying methods. With our guide we hope to increase understanding of the methods and hopefully clear up some of the confusion that exits regarding references and miss-citations of articles when citing methods.

By existing in the Matlab framework, the toolbox both makes it easier for users to customise their analysis to suit their experimental conditions and to make the analysis less tedious by being able to create script that can loop the analysis over an entire cohort of subjects and different conditions. We have also sought to thoroughly document the code of the toolbox. Both so users can see how the algorithms are implemented to increase transparency, but also it makes it easier to add new methods to the toolbox in the future.

This manuscript has been intended both as a guide to the toolbox and as an overview of the methods employed in the field of microstates. Therefore, if you have things that you feel are missing or find errors in either guide or the toolbox, please lets us know.

## Acknowledgements

We owe thanks to Frans Zdyb for helpful discussions and his preliminary work for a toolbox. We would also like to thank Scott Makeig for helpful discussions and comments for this manuscript. Finally, we thank Denis Brunet for helpful discussions regarding the TAAHC method.

The work was supported by the Novo Nordisk Foundation Interdisciplinary Synergy Program 2014, Biophysically adjusted state-informed cortex stimulation (BASICS) with grant number NNF14OC0011413, and by the Swiss National Science Foundation (100014_175875).

## A Optimised iteration scheme for modified K-means

The modified K-means algorithm is based on the loss function

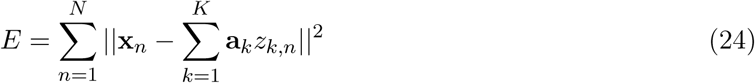

which should be minimised under the constraints

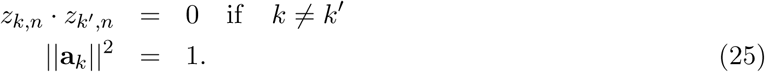

The first constraint ensures that a single microstate is active at any given time, while the second constraint resolves the scaling ambiguity between the **a** and the **z** variables. While closed form optimization is not feasible, we can consider two different iterative schemes.

We can follow a K-means approach as in Pascual-Marqui et al. (1995): Interchange hard assignment and **a** updates. In this case assume the assignments (each data point is assigned to a component *l*_*n*_) to be given. In this case we may first optimise the loss function w.r.t. the 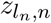 variables

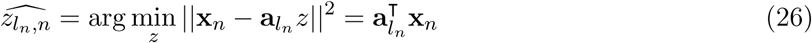

Substitute this activation measure in the loss function to obtain

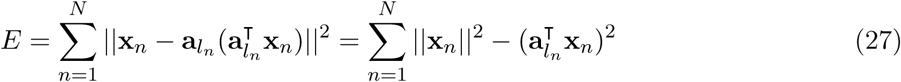

This loss function can now be minimised w.r.t. **a**_*k*_ under the unit length constraint, leading to the eigenvalue problem

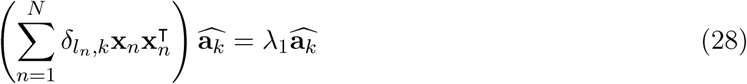

Where the delta function weighted sum 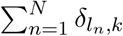 assures that only data points associated with the component, i.e., *l*_*n*_ = *k* are included.

Alternatively, we may consider an iterative scheme in which we first keep *z*_*k,n*_ fixed (i.e., both the assignments *l*_*n*_ and the activities 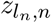) and optimise for the scalp maps **a**. This leads to

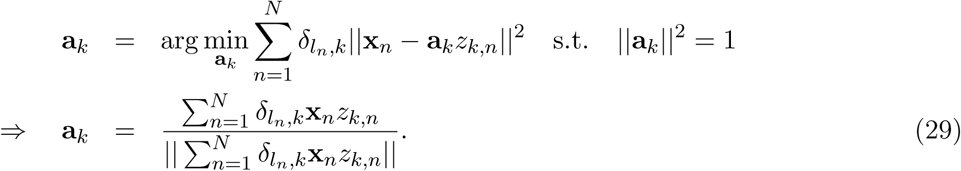

Next we obtain new values for *l*_*n*_ and *z*_*k,n*_ similar to the first approach.

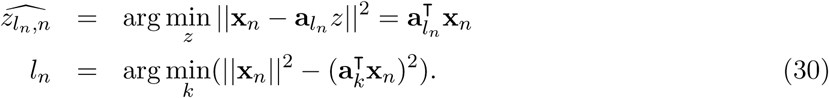

The two approaches lead to different updates for **a**. The second scheme is slightly faster than the first and turns out to produce an improved fit when empirically evaluated, see Figure 11.

**Figure 11:**
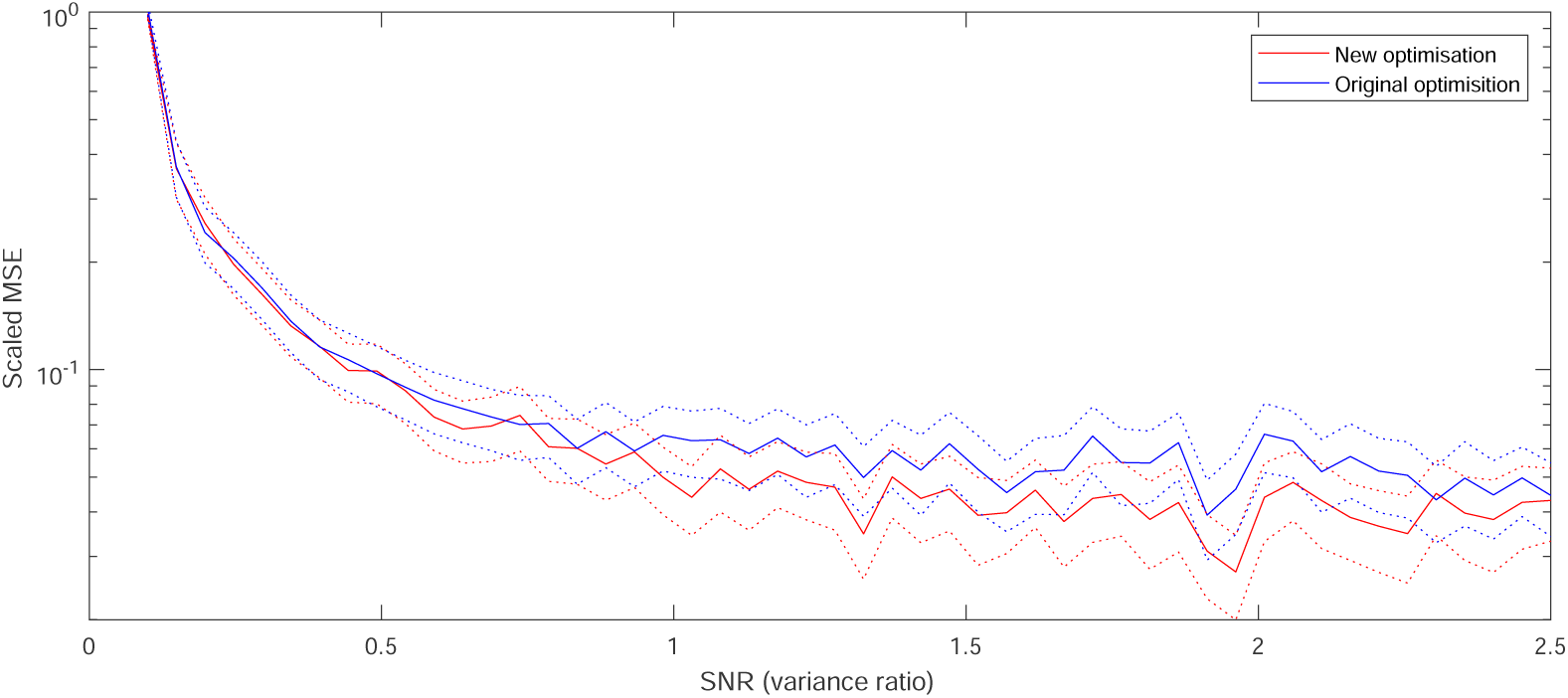
Simulation experiment to benchmark the two optimization procedures for modified K-means. Simulated data (*N* = 2000, *K* = 10, *C* = 30) and variable signal-to-noise rations (variance ratios) were analysed using *N*_its_ = 50 iterations (typical convergence took about 20 iterations). A scaled mean square error (MSE) measure is computed between the noise free signal, **X**, and the reconstructed signal, **AZ**^T^, as ‖**X**_0_ − **AZ**^T^‖^2^ */* ‖**X**_0_ **‖**^2^. For this simulated data, the new iteration scheme were on average ∼4 times faster than the original.

## MATLAB code to Tutorial

~~~
%% 3 Tutorial: EEG microstates analysis on spontaneous EEG Data
%
% This script executes the analysis steps of the Tutorial described in
% detail in section 3.3 to 3.8 of:
% Poulsen, A. T., Pedroni, A., Langer, N., & Hansen, L. K. (2018).
% Microstate EEGlab toolbox: An introductionary guide. bioRxiv.
%
% Authors:
% Andreas Trier Poulsen, atpo@dtu.dk
% Technical University of Denmark, DTU Compute, Cognitive systems.
%
% Andreas Pedroni, andreas.pedroni@uzh.ch
% University of Zurich, Psychologisches Institut, Methoden der
% Plastizitaetsforschung.
clear;clc;
% start EEGLAB to load all dependent paths
eeglab
%% set the path to the directory with the EEG files
% change this path to the folder where the EEG files are saved
EEGdir = ‘∼/EEGFiles/’;
% retrieve a list of all EEG Files in EEGdir
EEGFiles = dir([EEGdir ‘*.set’]);
%% 3.3 Data selection and aggregation
%% 3.3.1 Loading datasets in EEGLAB
for i=1:length(EEGFiles)
   EEG = pop_loadset(‘filename’,EEGFiles(i).name,’filepath’,EEGdir);
   [ALLEEG, EEG, CURRENTSET] = eeg_store(ALLEEG, EEG, 0);
   eeglab redraw % updates EEGLAB datasets
end
%% 3.3.2 Select data for microstate analysis
[EEG, ALLEEG] = pop_micro_selectdata(EEG, ALLEEG, ‘datatype’, ‘spontaneous’,… ‘avgref’, 1, …
   ‘normalise’, 0, …
   ‘MinPeakDist’, 10, …
   ‘Npeaks’, 1000, …
   ‘GFPthresh’, 1, …
   ‘dataset_idx’, 1:4);
   % store data in a new EEG structure
[ALLEEG EEG] = eeg_store(ALLEEG, EEG, CURRENTSET);
eeglab redraw % updates EEGLAB datasets
%% 3.4 Microstate segmentation
% select the “GFPpeak” dataset and make it the active set
[ALLEEG EEG CURRENTSET] = pop_newset(ALLEEG, EEG, 4,’retrieve’,5,’study’,0);
eeglab redraw
% Perform the microstate segmentation
EEG = pop_micro_segment(EEG, ‘algorithm’, ‘modkmeans’, …
   ‘sorting’, ‘Global explained variance’, …
   ‘Nmicrostates’, 2:8, …
   ‘verbose’, 1, …
   ‘normalise’, 0, …
   ‘Nrepetitions’, 50, …
   ‘max_iterations’, 1000, …
   ‘threshold’, 1e-06, …
   ‘fitmeas’, ‘CV’,… ‘optimised’,1);
[ALLEEG EEG] = eeg_store(ALLEEG, EEG, CURRENTSET);
%% 3.5 Review and select microstate segmentation
%% 3.5.1 Plot microstate prototype topographies
figure;MicroPlotTopo(EEG, ‘plot_range’, []);
%% 3.5.2 Select active number of microstates
EEG = pop_micro_selectNmicro(EEG);
[ALLEEG EEG] = eeg_store(ALLEEG, EEG, CURRENTSET);
%% Import microstate prototypes from other dataset to the datasets that should be
back-fitted
% note that dataset number 5 is the GFPpeaks dataset with the microstate
% prototypes
for i = 1:length(EEGFiles)
   fprintf(‘Importing prototypes and backfitting for dataset %i\n’,i)
   [ALLEEG EEG CURRENTSET] = pop_newset(ALLEEG, EEG, CURRENTSET,’retrieve’,i,’study’,0);
   EEG = pop_micro_import_proto(EEG, ALLEEG, 5);
%% 3.6 Back-fit microstates on EEG
EEG = pop_micro_fit(EEG, ‘polarity’, 0);
%% 3.7 Temporally smooth microstates labels
EEG = pop_micro_smooth(EEG, ‘label_type’, ‘backfit’, …
   ‘smooth_type’, ‘reject segments’, …
   ‘minTime’, 30, …
   ‘polarity’, 0);
%% 3.9 Calculate microstate statistics
EEG = pop_micro_stats(EEG, ‘label_type’, ‘backfit’, …
   ‘polarity’, 0);
[ALLEEG EEG] = eeg_store(ALLEEG, EEG, CURRENTSET);
end
%% 3.8 Illustrating microstate segmentation
% Plotting GFP of active microstates for the first 1500 ms for subject 1.
[ALLEEG EEG CURRENTSET] = pop_newset(ALLEEG, EEG, CURRENTSET,’retrieve’,1,’study’,0);
figure;MicroPlotSegments(EEG, ‘label_type’, ‘backfit’, …
   ‘plotsegnos’, ‘first’, ‘plot_time’, [4200 5700], ‘plottopos’, 1);
eeglab redraw
~~~

## Footnotes

1 But see: Koenig et al. (2011) for a Matlab implementation of microstates analyses for ERP data, or Milz (2016) for a Python implementation.

2 see section 2.1.2 for more information on polarity

3 This is also described in the Cartool help pages by Denis Brunet.

4 In the toolbox we have made polarity invariance optional, to enable customisation.

5 See section 2.2 for more information on GEV

6 Be careful not to confuse cross-validation with the CV criterion, which is a measure of fit.

7 We recommend reading up on cross-validation before attempting it, as it is possible to make unintentional mistakes, that breaks the division between training and test set.

8 The KL_nrm_ is the criterion implemented in stand-alone microstate software, CARTOOL.

9 Note that the article Lehmann and Skrandies (1980) is often cited for introducing GMD, and while it does introduce the concept of measuring topographical similarity between two microstate time samples, it does not introduce the equation. See e.g. Michel et al. (2009); Murray et al. (2008) for the equation and more information on GMD.

10 The original, raw-data is available at http://fcon_1000.projects.nitrc.org/indi/cmi_eeg/eeg.html

11 Available at https://github.com/amirrezaw/automagic

12 see https://sccn.ucsd.edu/wiki/Chapter_02:_Writing_EEGLAB_Scripts for a description on using this command

13 Given that the selected EEG maps at GFP peaks are temporally independent, the clustering algorithm ignores information about the order of their appearance, which is the reason that we do not apply temporal smoothing to the segmentation, in contrast to the microstate segmentation of event-related EEG.

14 See https://sccn.ucsd.edu/wiki/Chapter_01:_Loading_Data_in_EEGLAB for details on installing and starting.

15 In the case *ERP* has been selected, the epochs/ERPs from each dataset are either averaged into one grand average ERP or are concatenated in time (resulting in one long epoch).

16 The (T)AAHC algorithms are also polarity invariant and could be used. The K-means algorithm implemented in this toolbox, however, accounts for polarity because it uses the Euclidean distance as a similarity measure. K-means is therefore better suited for ERP analysis, where one might want to take polarity into account.

17 see section 2.5.1 for more information on the two options for temporally smoothing microstate labels.

